# Steroid signaling controls sex-specific development in an invertebrate

**DOI:** 10.1101/2023.12.22.573099

**Authors:** Lydia Grmai, Erin Jimenez, Ellen Baxter, Mark Van Doren

## Abstract

In vertebrate sexual development, two important steroid hormones, testosterone and estrogen, regulate the sex-specific development of many tissues. In contrast, invertebrates utilize a single steroid hormone, ecdysone, to regulate developmental timing in both sexes. However, here we show that in *Drosophila melanogaster*, sex-specific ecdysone (E) signaling controls important aspects of gonad sexual dimorphism. Rather than being regulated at the level of hormone production, hormone activity is regulated cell-autonomously through sex-specific hormone reception. Ecdysone receptor (EcR) expression is restricted to the developing ovary and is repressed in the testis at a time when ecdysone initiates ovary morphogenesis. Interestingly, EcR expression is regulated downstream of the sex determination factor Doublesex (Dsx), the founding member of the Dsx/Mab3 Related Transcription Factor (DMRT) family that regulates gonad development in all animals. E signaling is required for normal ovary development^1,2^, and ectopic activation of E signaling in the testis antagonized stem cell niche identity and feminized somatic support cells, which were transformed into follicle-like cells. This work demonstrates that invertebrates can also use steroid hormone signaling to control sex-specific development. Further, it may help explain recent work showing that vertebrate sexual development is surprisingly cell-autonomous. For example, chickens utilize testosterone and estrogen to control sex-specific development, but when they have a mixture of cells with male and female genotypes, the male cells develop as male and the female cells develop as female despite exposure to the same circulating hormones^3^. Sex-specific regulation of steroid hormone signaling may well underly such cell-autonomous sexual fate choices in vertebrates as it does in *Drosophila*.

---

Sexual dimorphism is a hallmark feature of metazoans and encompasses all discernible differences between males and females of the same species. Sex-specific gonad development is instructed by members of the Doublesex/Mab-3 Related Transcription Factor (DMRT) family in all animals for which their role has been characterized, including flies, nematodes, chickens, mice, and humans^4^. *DMRT1*-mutant male mice initiate testis development but the gonad then becomes feminized^5,6^, whereas in rabbits, loss of DMRT1 causes earlier sex reversal with ovary formation in XY embryos^7^. Humans that are XY, but also heterozygous for mutations in DMRT1, exhibit a range of phenotypes, including complete gonadal dysgenesis with sex reversal. Interestingly, these patients also have persistent Mullerian Ducts indicating an early problem in testis development (defects in Sertoli cells that produce anti-Mullerian Hormone)^8^.

The DMRT family was founded by the discovery of *doublesex (dsx)* in flies^9^. *Drosophila* sex determination leads to alternative splicing of *dsx* RNA and the production of either male or female isoforms of Dsx protein (Dsx^M^ or Dsx^F^), which share the same DNA binding domain but have opposite effects on target gene transcription. *dsx* controls all known aspects of sexual dimorphism in the somatic gonad, including important components of the germline stem cell niche (the “hub” in males and terminal filaments (TFs) and cap cells in females), and the somatic cells that support gametogenesis (cyst stem cells and cyst cells in males, and escort cells, follicle stem cells and follicle cells in females)(reviewed by^10^). The timing of gonad development is also sexually dimorphic, with the testis being formed by the end of embryogenesis (24 hrs after fertilization, AF), while ovary development occurs several days later in the mid-third larval instar (L3) stage (4 days AF).

In mammals, the steroid hormones testosterone and estrogen control many aspects of sex-specific development in a variety of tissues. In contrast, invertebrates like *Drosophila* utilize the same steroid hormone, ecdysone (E), in both sexes to regulate developmental timing and adult female reproduction^11^. The ecdysone receptor (EcR) is a nuclear hormone receptor, similar to the mammalian testosterone and estrogen receptors, that is activated by 20-hydroxyecdysone (20E). EcR-dependent transcriptional activation requires a number of co-factors that are also conserved in mammals, including the RXR ortholog Ultraspiracle (Usp) and the SRC-3 homolog Taiman (Tai). EcR target gene activation has been implicated in the establishment of the female gonad niche during development^1,2^ although its role in regulating sex-specific gonadogenesis has not been examined.

## *dsx* regulates ecdysone signaling in the gonad

We examined transcriptional activation via EcR in developing gonads using a transgenic reporter containing EcR binding sites (*EcRE-lacZ*)^12,13^ and immunostaining for the EcR target Broad (Br)^2^. In late larval ovaries, *EcRE-lacZ* produced high levels of β-gal throughout the apical cap, including TF cells (Fig. 1a). Similarly, Br was expressed strongly in all somatic cells of the late larval ovary, including the developing TFs^2^ (Extended Data Fig. 1a). In the larval testis, however, little to no *EcRE-lacZ* activity or Br expression was detected, including in the testis niche (Fig. 1b outline, Extended Data Fig. 1b). However, Br was detected in the testis terminal epithelium cells (Extended Data Fig. 1c), indicating that the testis environment receives 20E and supports ecdysone signaling in the cells that can respond. The Z1 isoform of Br is the most ecdysone-responsive isoform in the developing ovary^11^ (Extended Data Fig. 1d), and Br-Z1 was consistently absent from the male stem cell niche (Extended Data Fig. 1e).

**Figure 1.**
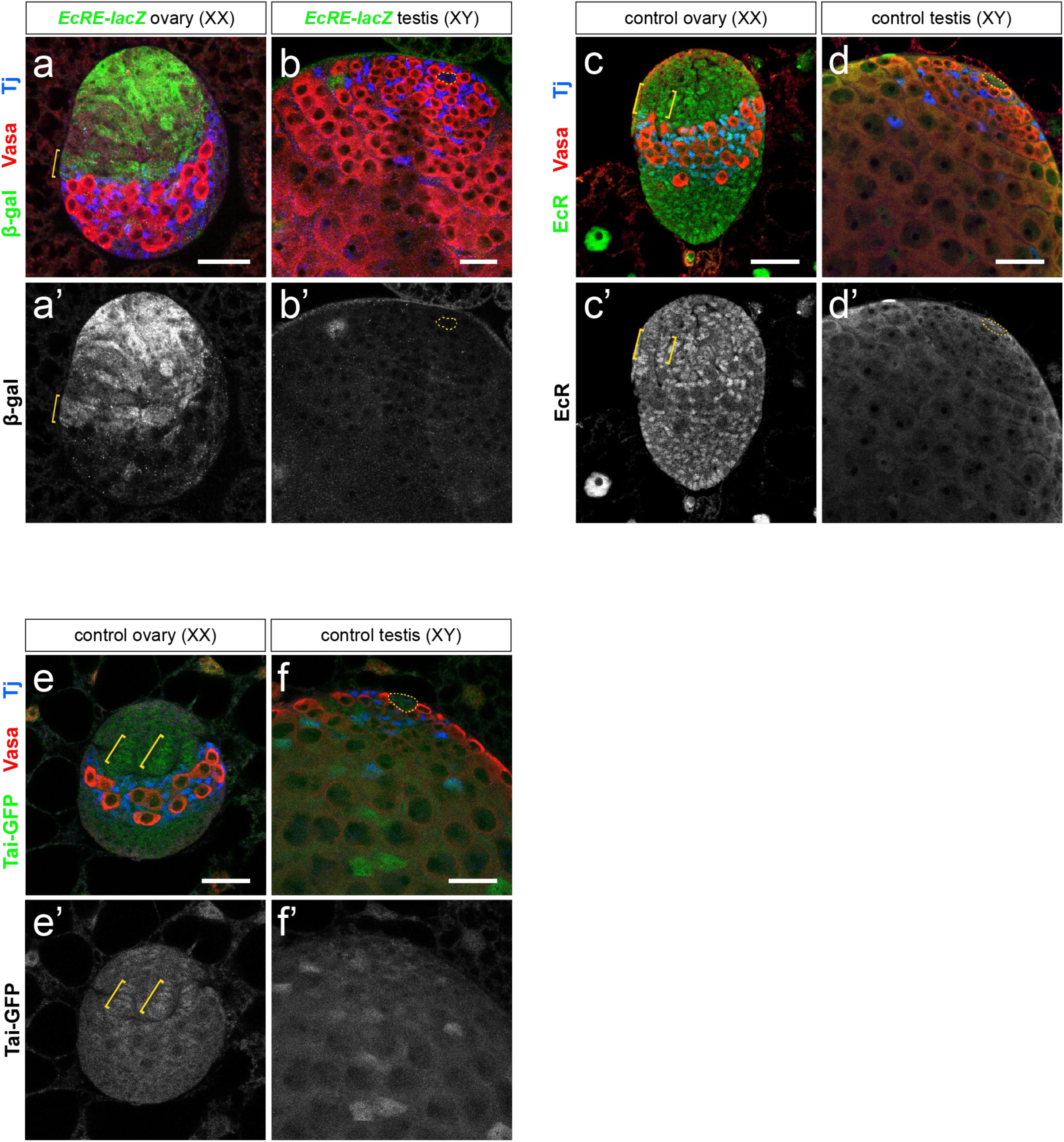
Ecdysone signaling activity is female-biased in the developing gonad. (a-b) Beta-galactosidase expression (green) from a transgenic *EcRE-lacZ*^25^ reporter is higher in the ovarian somatic gonad (a) than in the testicular somatic gonad (b). (c-d) EcR protein expression is higher in the ovary (c) than in the testis (d). (e-f) Expression of Taiman-GFP from a genomic duplication is detected in several somatic cell types of an L3 ovary (e), including the apical cap, terminal filaments, and intermingled cells. In an L3 testis (d), Tai is detected at low levels in the hub and CySCs and at higher levels in differentiating cyst cells. Vasa (red) labels the germline, Traffic jam (Tj, blue) labels somatic cells. Scale bars = 25 μm. Yellow brackets indicate TF cells; yellow dotted outline indicates the hub.

In mammals, sex-biased hormone activity is regulated at the level of hormone production: females produce higher levels of estrogen, while males produce higher levels of testosterone. In contrast, we found that a female bias in hormone levels could not account for the observed dimorphism in ecdysone activity. Supplying exogenous 20E, or a membrane-permeable EcR agonist (chromafenozide), did not activate E signaling in testes but was able to induce premature EcR transcriptional activity in younger L3 ovaries (Extended Data Fig. 2a-f). Ecdysone is converted to its active form, 20E, by the P450 enzyme Shade^27^ (Shd). We found that depletion of *shd* from the ovary did not reproduce female niche defects that are seen upon blocking EcR activity^2^ (Extended Data Fig. 2g-h). We conclude that the female bias in E signaling is not due to sex-specific production or availability of 20E.

We previously conducted a collaborative genome-wide search for putative Dsx targets using genomic and computational approaches^14^. We identified numerous sequences across the *EcR* locus with high similarity to the Dsx consensus binding sequence, many of which were conserved in other *Drosophila* species and could be bound by Dsx based on DamID-seq^14^. We therefore explored whether sex differences in *EcR* expression might account for sexually dimorphic E signaling in the gonad. EcR was expressed at high levels in the female somatic gonad (Fig. 1c) and at very low levels in the larval testis (Fig. 1d). We also saw expression of EcR in the terminal epithelium of the testis (Extended Data Fig. 1h, k) which were the only cells of the testis that expressed the *EcR* target BrC (Extended Data Fig. 1c). An EcR co-factor, Taiman (Tai), that mediates ecdysone signaling in the adult ovary^15^ was expressed in the somatic gonad of both larval ovaries and testes (Fig. 1e-f).

To test whether EcR expression and activity are regulated downstream of *dsx*, we assessed EcR and Br levels in *dsx-*mutant gonads. We generated female animals heterozygous for the *dsx^D^* allele^16^, which cannot be spliced into the *dsx^F^*isoform. Thus, XX *dsx^D^/+* animals express both Dsx^M^ and Dsx^F^, which antagonize one another^16^. In the gonad, this leads to stochastic establishment of male or female niche structures (hubs or TFs)^17^. Importantly, all other aspects of the female sex determination pathway besides *dsx* are present in these animals, making this a good test of *dsx* function. We found that XX *dsx^D^/+* gonads exhibited lower levels of EcR protein (Fig. 2a-d) and *EcR* mRNA (Extended Data Fig. 3a-c) compared with control ovaries. While Br-Z1 was predictably female-specific in control gonads (Fig. 2e-f), in *dsx^D^/+* gonads Br-Z1 expression in the niche correlated with niche identity; *dsx^D^/+* gonads with a hub expressed very low levels of Br-Z1 (Fig. 2g) while *dsx^D^/+* gonads with TFs expressed higher levels of Br-Z1 (Fig. 2h), supporting the idea that high ecdysone signaling promotes female niche development.

**Figure 2.**
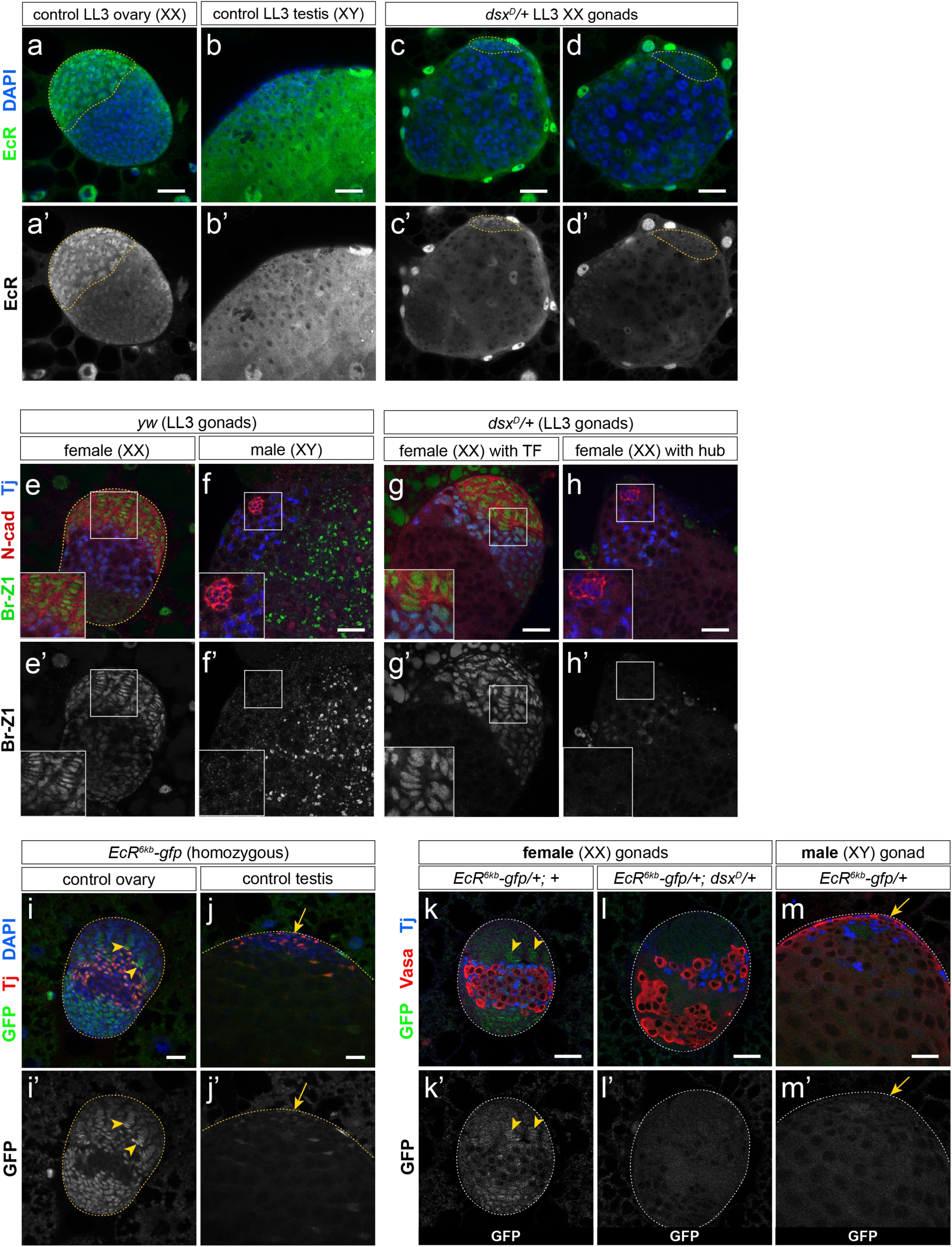
EcR activity is regulated by Dsx in the developing somatic gonad. (a-c) EcR protein levels are higher in an L3 ovary (a, green) than in an L3 testis (b). EcR expression is diminished in XX *dsx*-mutant gonads (*dsx^D^/+*, c-d). DAPI (blue) labels nuclei. (e-h) Br-Z1, an EcR target, is expressed in TFs of a control ovary (e) and is absent from hub cells in a control testis (f). Br-Z1 expression is high in an L3 XX *dsx^D^/+* TF structure (g) and low in a hub structure (h). N-cad labels the gonad niche (TF or hub cells) (i-j) Expression of a 6.3-kb intronic EcR enhancer (*EcR^6kb^-gfp*) in larval gonads. Gonads shown in (i) and (j) contain two copies of enhancer construct (homozygous). *EcR^6kb^-gfp* is expressed throughout the somatic gonad of an LL3 ovary (h, arrowheads) and is expressed at very low levels in the hub (arrowhead) and CySCs of an L3 testis (j). (k-l) *EcR^6kb^-gfp* expression is greatly reduced in *dsx*-mutant (*EcR^6kb^-gfp/+; dsx^D^/+*) gonads (l) compared with control (*EcR^6kb^-gfp/+; +*) ovaries (k). Heterozygous expression of *EcR^6kb^-gfp* in a control testis is shown in (m). In k-m, DAPI (blue) labels nuclei and Tj (red) labels somatic cells. Endogenous GFP signal (without GFP antibody staining) is shown in green. In i-m, arrowheads indicate TF cells and arrows indicate the hub. Scale bars = 25 μm.

To evaluate whether *dsx* is a transcriptional regulator of *EcR*, we generated a transgenic reporter, *EcR^6kb^-gfp*, that expresses GFP under control of a 6.3-kb intronic enhancer element in the *EcR* locus and is expressed in the somatic gonad. *EcR^6kb^-gfp* was expressed in somatic cells of the larval ovary, while very low levels were observed in the larval testis (Fig. 2i, j). Interestingly, *EcR^6kb^-gfp* activity was lower in XX *dsx^D^/+* gonads compared with control XX gonads (Fig. 2k-m; Extended Data Fig. 3d-f) indicating that the effect of *dsx* on *EcR* expression is transcriptional.

## EcR antagonizes testis development

Since EcR activity was female-biased during gonad development, we investigated whether the requirement for *EcR* in gonad niche development is sexually dimorphic. EcR activation by 20E is required for development of the female gonad stem cell niche^1,2^. To investigate the role of EcR in gonad niche development, we blocked EcR activation in the somatic gonad by over-expressing a dominant-negative form of *EcR-A*^18^ (*EcR^DN^*) using the somatic gonad driver *traffic jam (tj)-GAL4* (*tj-GAL4; UAS-EcR^DN^*, abbreviated as *tj>EcR^DN^*). EcR acts as a repressor in the absence of 20E^19^, which is required to prevent premature ovary development^2^, preventing the use of *EcR* null alleles for this experiment. However, expression of *EcR^DN^* specifically blocks the activator function of EcR^18^. As previously observed^2^, we found that *EcR^DN^*expression in the somatic gonad prevented the differentiation of female niche structures (TFs, Fig. 3a, b). In contrast, transcriptional activation via EcR was not required for establishment of the male gonad niche, as testes expressing *EcR^DN^* did not contain fewer hub cells than control testes. In fact, *tj>EcR^DN^* testes showed a slight increase in hub cell number and hub volume compared with controls (Extended Data Fig. 4).

**Figure 3.**
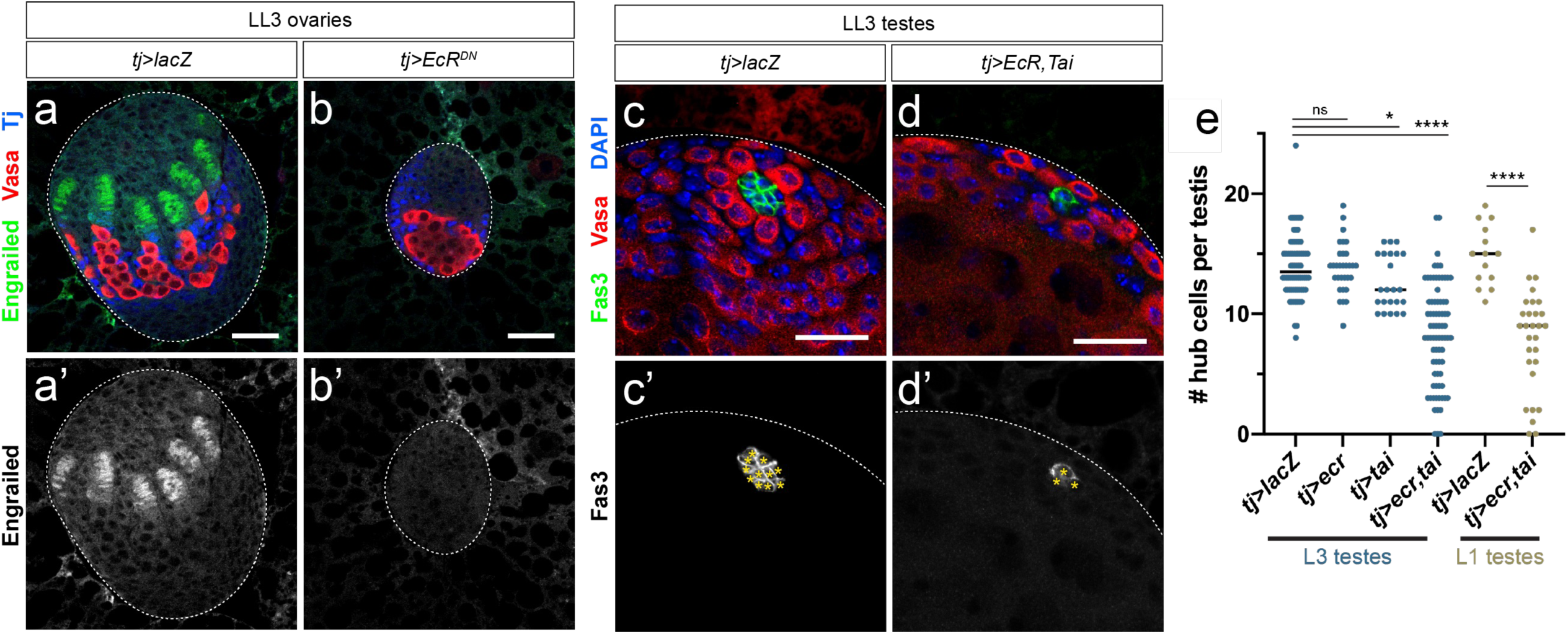
EcR signaling is deleterious to male somatic gonad establishment. (a) TFs are established in a control LL3 ovary (*tj>lacZ*). Engrailed (green) labels TF cells. (b) Blocking EcR activation in the somatic gonad (*tj>EcR^DN^*) prevents the establishment of terminal filaments in an L3 ovary. (c) In a control L3 testis, Fas3 (green) labels intact hub cells. (d) Activation of EcR by co-expressing EcR and Tai (*tj>EcR,tai*) in the male somatic gonad produces smaller hubs with visibly fewer cells. Fas3 (green) labels hub cells. (g) Quantification of hub cells per LL3 (blue dots) or L1 (yellow dots) testis in various genotypes. For statistical significance here and in subsequent figures, asterisks indicate statistical significance by Student’s t-test as follows: *p<0.05; **p<0.01; ***p<0.001; ****p<0.0001. Scale bars = 25 μm.

**Figure 4.**
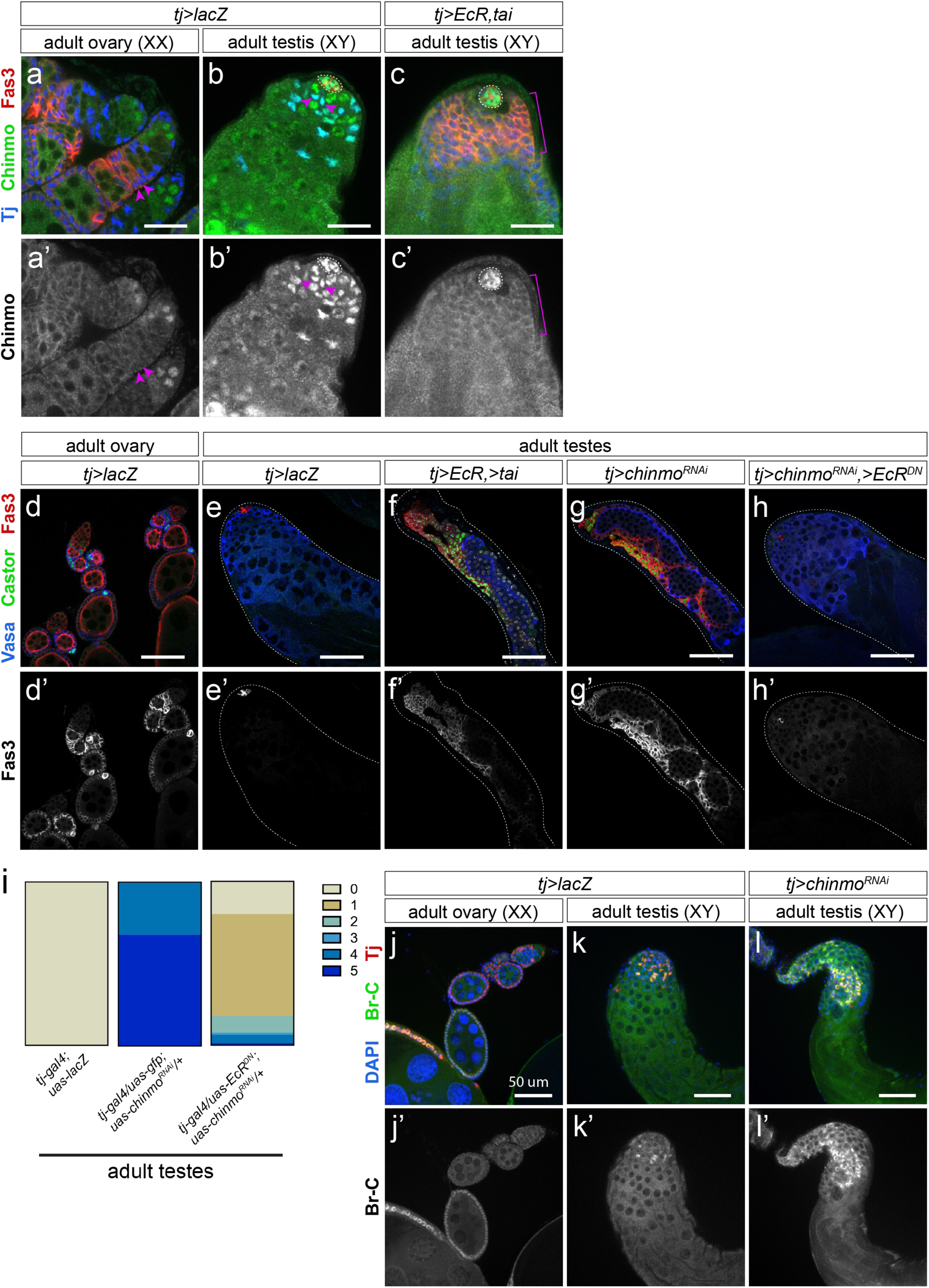
EcR activation in the testis causes adult somatic sex transformation. (a-c) Chinmo expression in adult gonads and with EcR activation in the testis. In an adult ovary (a), Chinmo protein (green) is absent from the female somatic gonad (arrowheads). Chinmo is expressed in early female germ cells as previously observed^3^. (b) In an adult testis, Chinmo is detectable in the hub and CySCs. (c) Chinmo expression is dramatically reduced upon EcR activation in the adult testis. (d-h) Expression of follicle cell markers Fas3 (red) and Castor (green) in adult gonads. Adult ovaries (d) show expression of Fas3 (red) and Castor (green) in early follicle cells, while in a control adult testis (e) Castor and Fas3 are absent from the cyst cell lineage. In adult testes, as seen in previous figures, Fas3 is expressed in hub cells. EcR activation in the testis (f) causes somatic feminization in the adult testis, characterized by the presence of Fas3- and Castor-positive somatic aggregates. Depletion of *chinmo* in the male somatic gonad (g) also causes somatic feminization as previously observed^3^. Blocking EcR activation by over-expressing *EcR^DN^* (h) suppresses somatic feminization in the absence of Chinmo. (i) Quantification of feminization rescue in *tj>chinmo^RNAi^* testes by blocking EcR activation. An abbreviated description of phenotypic categories: 0=no feminization or germline defects; 1=germline defects but no feminization; 2=mild feminization; 3=moderate feminization; 4=severe feminization; 5=acellular testes. Feminization and acellularity are indicated on the graph in blue. More complete descriptions and visual examples of each phenotypic category can be found in Extended Data Fig. 9. (j-l) Expression of Br-C (green) in the adult testis upon *chinmo* depletion. Br-C is expressed in follicle cells during mid-oogenesis in the adult ovary (j). Br-C is expressed at moderate levels in early cyst cells but is absent from differentiating cyst cells in an adult testis (k). In a *tj>chinmo^RNAi^*testis (l), Br-C is expressed in later cyst cells distal from the hub (hub and testis apex are out of frame in l and l’). Scale bars = 50 μm.

To test whether activating EcR in the developing testis is deleterious to male niche development, we first identified the rate-limiting components for E signaling in the larval testis. Over-expression of EcR alone was not sufficient to activate signaling, as determined by quantifying Br expression (Extended Data Fig. 5a-e). However, co-expression *EcR* along with its co-factor *taiman* (*tj>EcR,tai*) in the male somatic gonad led to a significant increase in Br expression (Extended Data Fig. 5a-e) indicating we could now ectopically activate E signaling in the testis. This led to a significant decrease in larval hub cell number compared with control testes (Fig. 3c-e) which was apparent as early as the L1 stage (24-48 hrs AF, Fig. 3e and Extended Data Fig. 6a-c). Hub cell loss continued beyond larval development, as some *tj>EcR,tai* pupal testes lacked hubs and, consistent with this, also lost the germline and somatic support cells dependent on the male niche (Extended Data Fig. 6d-e). While EcR activation caused a loss of male niche identity, we did not find a transformation to female niche identity, as occurs in loss of *dsx* function^17^. Expression of the female TF marker Engrailed increased in male somatic cells upon activation of E signaling (Extended Data Fig. 7) but no evidence of TF formation was observed.

## EcR activity feminizes the adult testis

While activation of E signaling was not sufficient to sex-transform the gonad niche, we observed a striking male-to-female transformation of somatic support cells in the testis upon co-expression of EcR and Tai. Support cells in the adult testis include the cyst cells that surround each developing spermatogenic cyst and the cyst stem cells that produce them. In the ovary, somatic support cells include escort cells that nurture germ cells during the earliest stages of differentiation and epithelial follicle cells, made by follicle stem cells, that support maturing oocytes. The transcription factor Chronologically inappropriate morphogenesis (Chinmo) is normally expressed in hub cells and early cyst cells of L3 and adult testes, while in females it is expressed in germ cells but is absent from the somatic gonad at both stages^20–22^ (Fig. 4a, b and Extended Data Fig. 8a-b). Co-expression of EcR and Tai caused a loss of Chinmo expression in cyst cells but not hub cells (Fig. 4c). Interestingly, this was accompanied by a gain in expression of the follicle cell markers Fas3 and Castor (Fig. 4d-f), which also occurs upon Chinmo depletion from the testis^21,22^ (Fig. 4g). Repression of *chinmo* and activation of *br* by ecdysone signaling has also been observed in developing neuroblasts and in larval wing imaginal discs^23,24^ and has been proposed to regulate many aspects of development^25^. To investigate further the mutually repressive relationship between EcR and *chinmo* in the somatic gonad, we examined whether transcriptional activation via EcR was required for the male-to-female transformation observed upon loss of *chinmo*. Indeed, interfering with EcR activation by expressing *EcR^DN^* blocked the follicle cell-like transformation observed with loss of *chinmo* (Fig. 4g-h). Follicle-like cells were present in 92.3% of *tj>chinmo^RNAi^* testes, but they appeared in only 7.3% of *tj>EcR^DN^; >chinmo^RNAi^*testes (Fig. 4i). In addition, depletion of *chinmo* in the testis led to induction of the EcR target Br in the transformed follicle-like cells (Fig. 4j-l). These data are consistent with a model where EcR/Br and *chinmo* exhibit mutually repressive interactions in the somatic gonad.

## Discussion

Taken together, our data demonstrate that E signaling can be used to control sex-specific development in an invertebrate. This does not require sex-specific E production and, indeed, sufficient E is produced in both sexes to activate signaling in cells competent to respond to the hormone, such as the testis terminal epithelium (Extended Data Fig. 1a, c). We demonstrate that at the time that E signaling activates ovary development, E signaling is repressed in most somatic cells of the larval testis, including the hub and somatic support cells. Further, our work shows that this repression is necessary for proper testis development: activation of E signaling causes a loss of male niche (hub) identity and a transformation of adult somatic support cells toward a female follicle cell-like identity. We also show that repression of E signaling in the testis is due, in part, to *dsx*-dependent regulation of *EcR* expression. Therefore, invertebrates like *Drosophila* that utilize the steroid hormone E to regulate development in both sexes can also employ this same hormone to regulate sexual development via sex-specific regulation of its receptor.

We show that somatic gonad development and maintenance feature mutually exclusive expression of the transcription factors Chinmo and Br. Repression of *chinmo* by EcR and Br has been observed in other developmental contexts^23–25^, presenting this as a common regulatory mechanism. Thus, sex-specific EcR expression may well be used to control sexual dimorphism in other tissues such as the nervous system. It is notable that upon EcR hyperactivation in the adult testis, somatic cells appear transformed to a female follicle cell-like identity, whereas hub cells are lost but do not undergo a similar transformation toward a female niche identity. Male to female niche transformation is observed in *dsx* mutants^17^, indicating that there must be additional regulatory mechanisms downstream of *dsx*, in addition to EcR, that promote sex-specific niche development and maintenance. Candidates for this include *fruitless,* which encodes a transcription factor involved in hub maintenance during larval stages^26^, and *bric a brac 1* and *2 (bab1/2)*, which are important for TF cell specification^27,28^. Interestingly, Fruitless, Bab1/2, Br, and Chinmo all encode transcription factors with BTB (Broad, Tramtrack, Bric a brac) domains that promote homotypic and heterotypic protein interactions. Further, they are all either predicted or known Dsx target genes^1417,26,29^. Thus, the regulation of these factors by Dsx and EcR, in addition to their potential physical associations, could allow for a particularly rich network of regulatory interactions controlling sexual dimorphism in the gonad and other tissues.

The autonomous regulation of ecdysone signaling we report in *Drosophila* gonads may provide a model for how cell-autonomous sexual identity is regulated in vertebrates. Birds, like mammals, utilize estrogen and testosterone to control development of sexually dimorphic characteristics. Indeed, manipulating the activity of aromatase, which converts testosterone to estrogen, can induce sex-reversal during chicken development^30^. However, in birds with a mixture of cells with male and female sex chromosome genotypes, the cells with a male genotype develop as male and the cells with a female genotype develop as female, despite the fact that these animals have a single circulating level of estrogen and testosterone^3,31^. Further, male chickens heterozygous for a non-functional allele of *DMRT1* develop an ovary rather than testes, but all other tissues analyzed are phenotypically male^32^. These data clearly indicate that even when estrogen and testosterone are used to control sex-specific development, the responding tissues can cell-autonomously differ in how they respond to these hormones. The simplest explanation for this is that these tissues autonomously regulate their response to these steroid hormones according to their sex, similarly to how *Drosophila* cells regulate sex-specific response to ecdysone signaling in the gonads.

## Acknowledgments

We thank the Bloomington *Drosophila* Stock Center (Bloomington, IN, USA), the *Vienna* Drosophila Resource Center (Vienna, Austria), and the Developmental Systems Hybridoma Bank (Iowa City, IA, USA) for making available fly stocks and antibodies. We are also grateful to Van Doren Lab members and Dr. Shyama Nandakumar for project and manuscript feedback. This work was funded by NIH R01GM113001 (MVD).

## Author contributions

Conceptualization: L.G., E.J. and M.V.D.; Methodology: L.G., E.J., E.B., and M.V.D.; Investigation: L.G., E.J., and E.B.; Formal analysis: L.G. and E.J.; Validation: L.G., E.J. ad E.B.; Resources: M.V.D.; Writing – original draft: L.G.; Writing – review and editing: L.G. and M.V.D.; Supervision: L.G. and M.V.D.

## Methods

### Fly stocks and husbandry

The following flies were used and are described in FlyBase: *Oregon^R^*, *dsx^1^*, *Df(3R)dsx^3^*, *dsx^D^*, *msl3*-*gfp*, *tj-gal4 (P{GawB}NP1624)*, *EcRE-lacZ, UAS-lacZ*, *UAS-EcR.C*, *UAS-Tai*, *UAS-EcR-A^W650A^* (referred to as *UAS-EcR^DN^*in text), *UAS-chinmo^RNAi^* (HMS00037) tai-GFP. *EcR^6kb^-gfp* enhancer line was generated in this study; see below for cloning details.

### Antibodies

The following antibodies were used: rabbit anti-Vasa (1:10,000; gift of Ruth Lehmann, MIT/Whitehead Institute, Cambridge, MA, USA), guinea pig anti-Tj (1:1000), mouse anti-Fas3 (1:100; DSHB), rat anti-N-cadherin (1:20; DSHB), mouse anti-EcR (Ag10.2; 1:20; DSHB), mouse anti-EcR-A (15G1a; 1:20; DSHB), mouse anti-EcR-B1 (AD4.4; 1:20, DSHB), mouse anti-Broad-core (25E9.D7; 1:40; DSHB), mouse anti-Broad-Z1 (1:50; DSHB), mouse anti-β-galactosidase (40-1a; 1:40; DSHB), mouse anti-Engrailed (1:10; DSHB), rat anti-Chinmo (1:5,000; gift of N. Sokol, formerly Indiana University, Bloomington, IN), rabbit anti-Castor (gift of W. Odenwald, NIH, Bethesda, MD). Cross-adsorbed secondary antisera (Invitrogen) were raised in goat and diluted to 4 μg/mL for staining.

### Larval hormone feeding experiments

Larval 20-hydroxyecdysone (20E) and chromafenozide (CF) feeding experiments were performed as follows. First, embryos were collected for 4 hours on apple juice agar petri dishes supplemented with regular yeast paste. 21 hours after the start of embryo collection, larvae that hatched prematurely were removed. First instar larvae that hatched over the following 4 hours were then transferred to a fresh apple juice plate supplemented with regular yeast paste. Newly ecdysed third instar larvae were transferred to a fresh apple juice plate supplemented with either control yeast paste (dry yeast mixed with H_2_O lacking hormone), yeast paste plus 20E (dry yeast mixed with 1 mM 20E in H_2_O), or yeast paste plus 1 mM CF (dry yeast mixed with 1 mM CF in H_2_O). Wandering third instar larvae were dissected 48 hours later for gonad staining and imaging.

### Immunofluorescence

Testes and ovaries were dissected in PBS and fixed in 5% formaldehyde (diluted in PBS) for 20 minutes at room temperature (RT). Fixed tissue was washed twice for 10 minutes each in PBTx (PBS + 0.1% Triton X-100). Blocking was performed for 30 minutes or overnight in BBTx (PBS + 0.1% Triton X-100 + 1% BSA). Primary antibodies were diluted in BBTx and incubated overnight at 4° C. Primary antibodies were washed off twice for 10 minutes each in BBTx. Secondary antibodies were diluted in BBTx, incubated for 2-3 hours at RT in the dark, and washed off in PBTx. Samples were then incubated in 1 μg/mL DAPI for 10 minutes at RT in the dark, and finally washed twice in PBTx. HCR-FISH was performed according to manufacturer’s protocol (Molecular Instruments, Inc.) and *EcR* hybridization probes were generated commercially. Tissue mounting medium was supplemented with DABCO anti-fade reagent (*company*) prior to confocal analysis. Confocal images were captured using a Zeiss LSM 710 confocal microscope using 20x, 40x, or 63x objective lens.

### ImageJ quantifications

For quantification of *EcR^6kb^-gfp* gonad expression shown in Fig. 2, total GFP intensity from a representative Z-slice was quantified for each gonad, then normalized to total DAPI intensity within the same area.

### Generating EcR^6kb^-GFP constructs

Cloning was performed using a modified version of pStinger^46^ that replaces P-elements with *attB* site to facilitate ΦC31-mediated, site-directed genome integration. The MCS from *pStinger* (Barolo et al. 2000) was excised and ligated into *pARE-GFP^nls^* (see Chatterjee & Bohmann 2012 for vector map) to generate *pStinger-attB*. The *hsp70* minimal promoter sequence was removed (since the 6.3kb putative enhancer contains a core promoter) to generate *pStinger-attB*. This vector was used as the starting point to generate *EcR^6kb^-GFP*. The 6.3-kb putative enhancer region was amplified from a BAC construct containing part of the *EcR* locus (*BACR08A11*) using the primers listed below, then digested with SphI/AgeI and ligated into *pStinger-attB* using the NEBuilder HiFi DNA cloning assembly kit (New England Biosciences, Inc.). Positive clones were screened by PCR and sequence integrity was verified using Sanger sequencing. Plasmid was injected into *Drosophila* embryos and stable lines were generated via BestGene, Inc.

### Primers

EcR6kb_sphI_ageI_fwd: tagtgctactgcatagca EcR6kb+sph1_ageI_rev: ctatgcagccgccatata

---

**Extended Data Figure 1.**
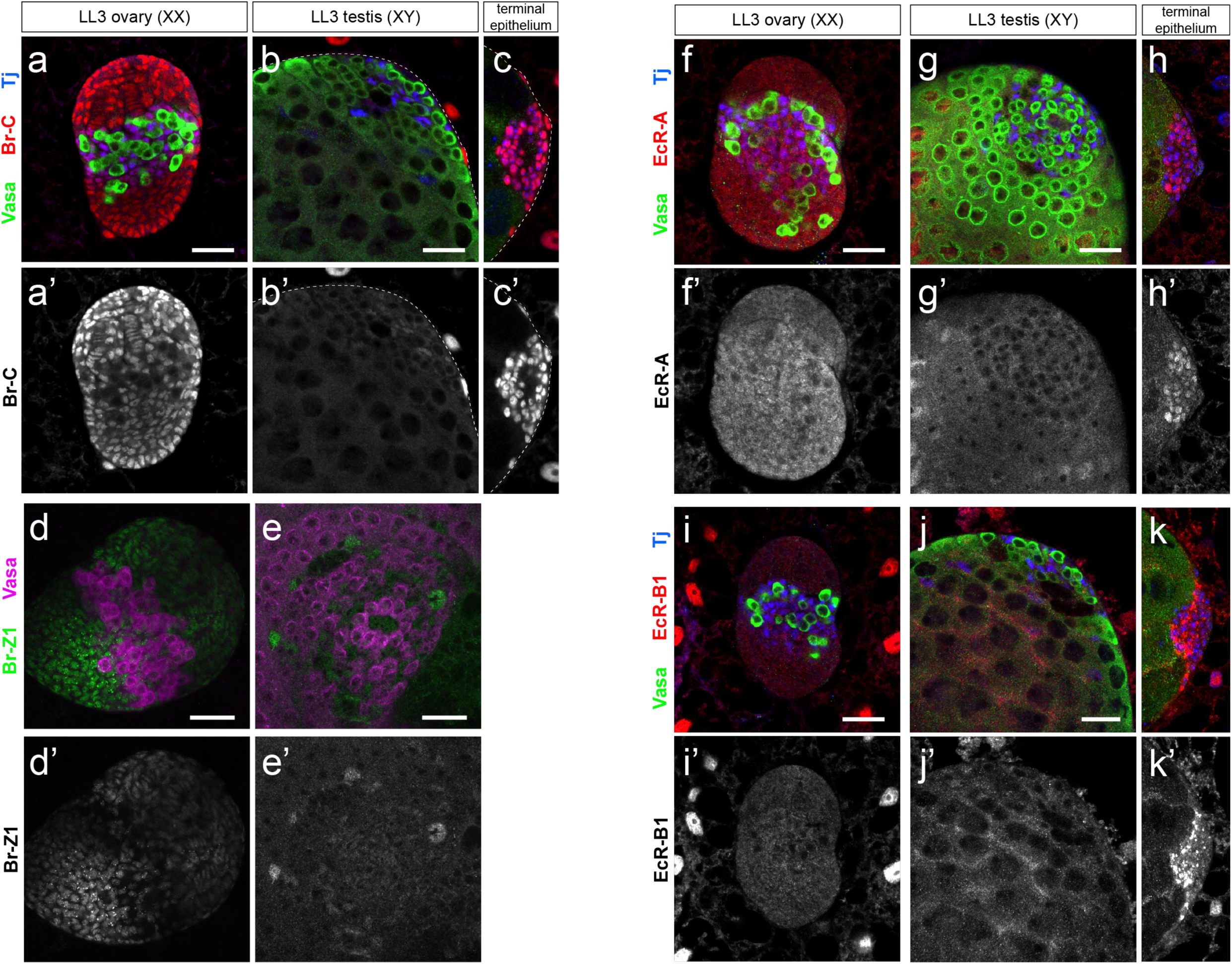
Expression of EcR and Br isoforms in the developing gonad. (a-b) Expression of the EcR target Br-C in an LL3 ovary (a) and testis (b). Br-C expression in terminal epithelium cells of a larval testis is shown in (c). Vasa (green) labels the germline and Tj (blue) labels somatic cells. (d-e) Expression of Br-Z1 isoform in an LL3 ovary (d) and testis (e). Vasa (magenta) labels the germline. (f-k) Expression of EcR-A (f-h) and EcR-B2 isoforms in the LL3 ovary (f, i), testis (g, j) and testicular terminal epithelium cells (h, k). Vasa (green) labels the germline and Tj labels somatic cells. Scale bars = 25 μm.

**Extended Data Figure 2.**
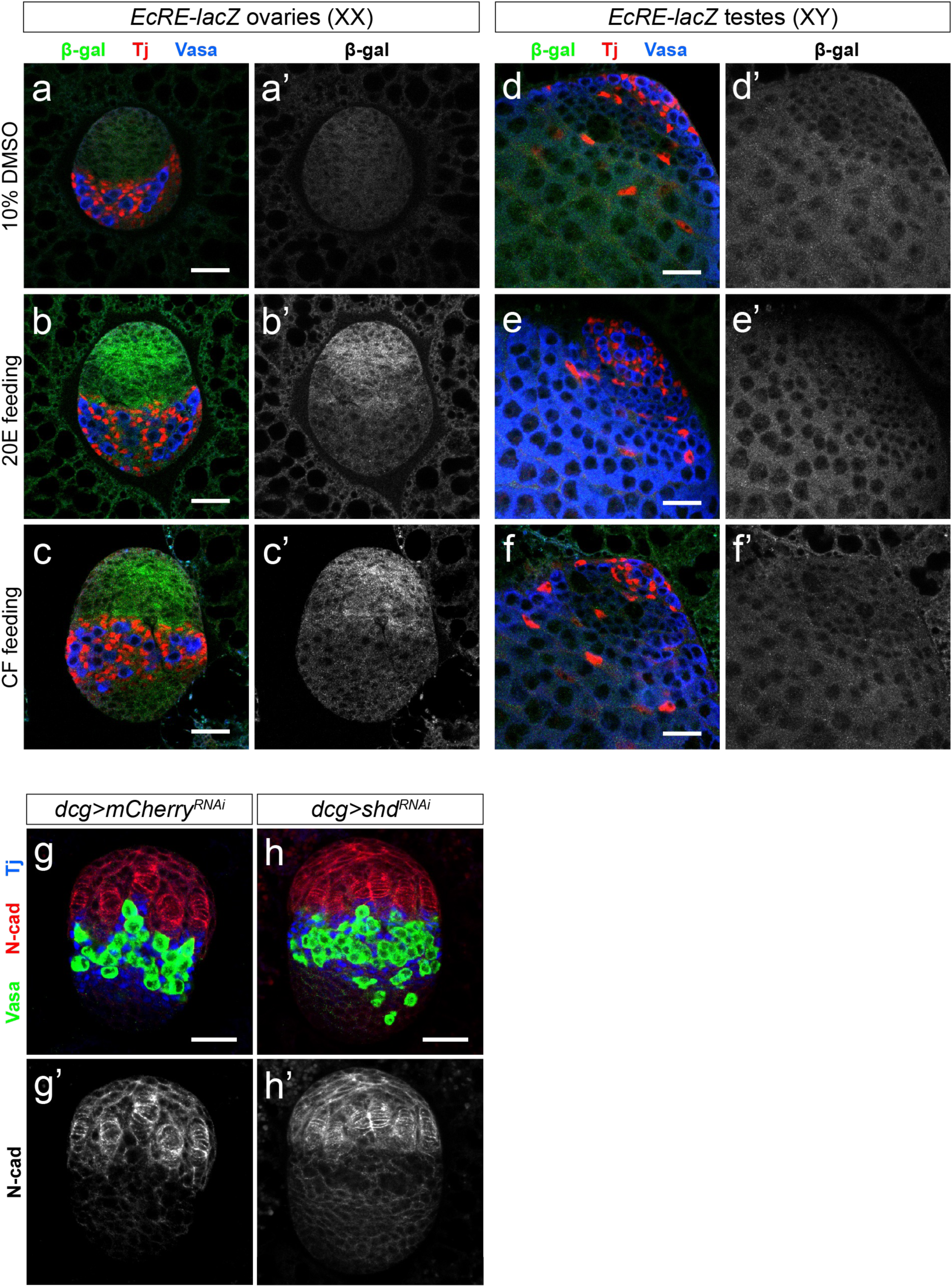
(a-f) Expression of *EcRE-lacZ* reporter in L3 ovaries (a-c) or L3 testes (d-f) upon feeding with 10% DMSO (vehicle, a and d), 20-hydroxyecdysone (20E, b and e), or chromafenozide (CF, c and f). (g-h) Depletion of *shd* in the larval ovary does not visibly impair ovary development. Note that *dcg-gal4* used in this experiment is expressed in both the ovary and surrounding fat body. Vasa (green) labels the germline, N-cad (red) labels TF cells, and Tj (blue) labels intermingled somatic cells. Scale bars = 25 μm.

**Extended Data Figure 3.**
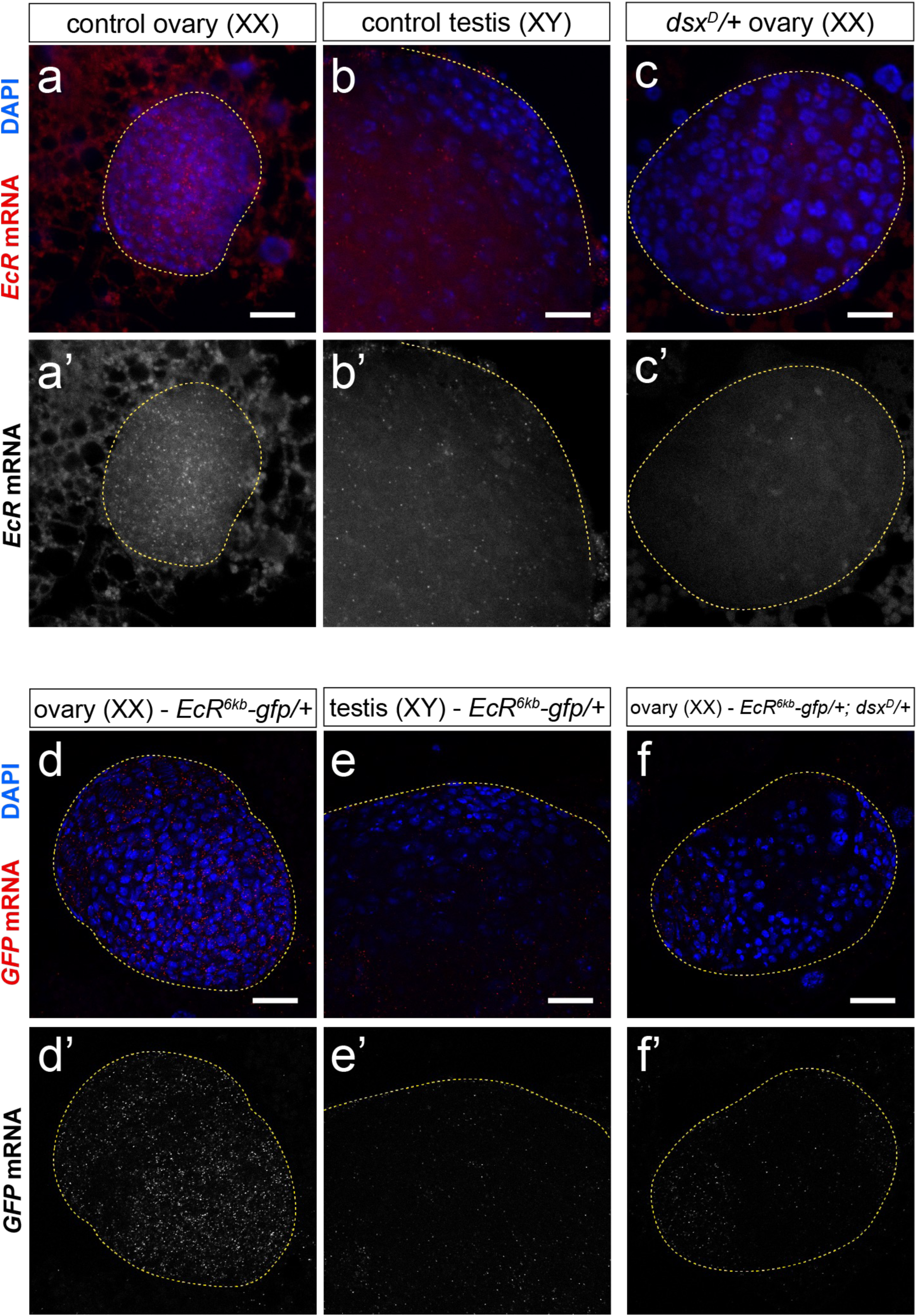
(a-c) Detection of *EcR* mRNA (red) in LL3 gonads by HCR-FISH. A *dsx*-mutant (*dsx^D^/+*) gonad (c) shows lower *EcR* mRNA abundance than control ovaries (a). *EcR* mRNA in an LL3 testis is shown in (b). (d-f) Detection of *GFP* mRNA (red) in LL3 gonads by HCR-FISH. *dsx^D^/+* gonad (f) contains higher *GFP* mRNA abundance than control ovaries (d). Scale bars = 25 μm.

**Extended Data Figure 4.**
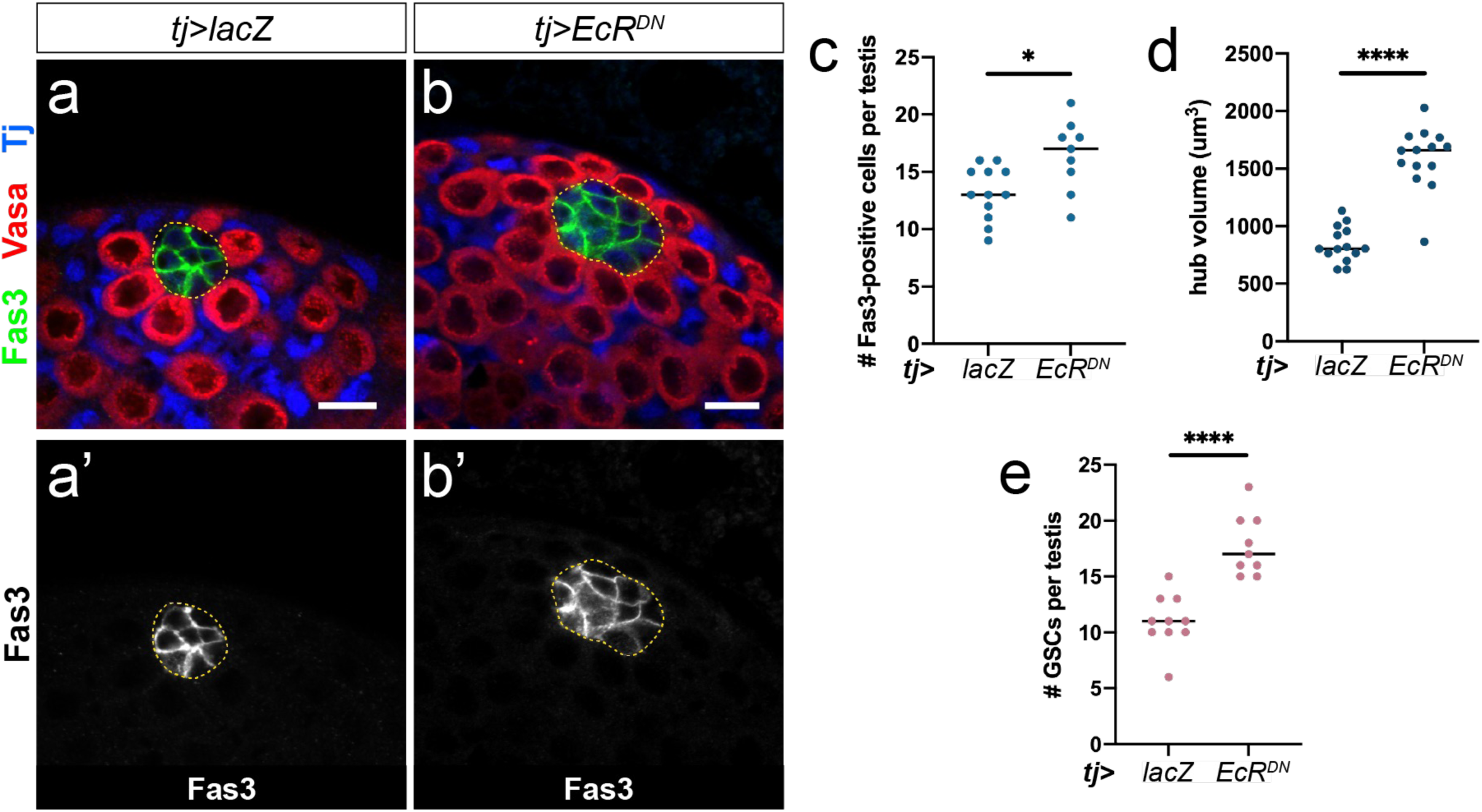
(a-b) Neither hub specification nor hub maintenance are compromised by blocking EcR activity (*tj>EcR-^DN^*). Fas3 (green) labels hub cells, Vasa (red) labels the germline, and Tj (blue) labels somatic cells. (c-e) Quantification of hub cell number (c), hub volume (d), and germline stem cell (GSC) number (e) upon blocking EcR activation in the somatic gonad. Scale bars = 10μm.

**Extended Data Figure 5.**
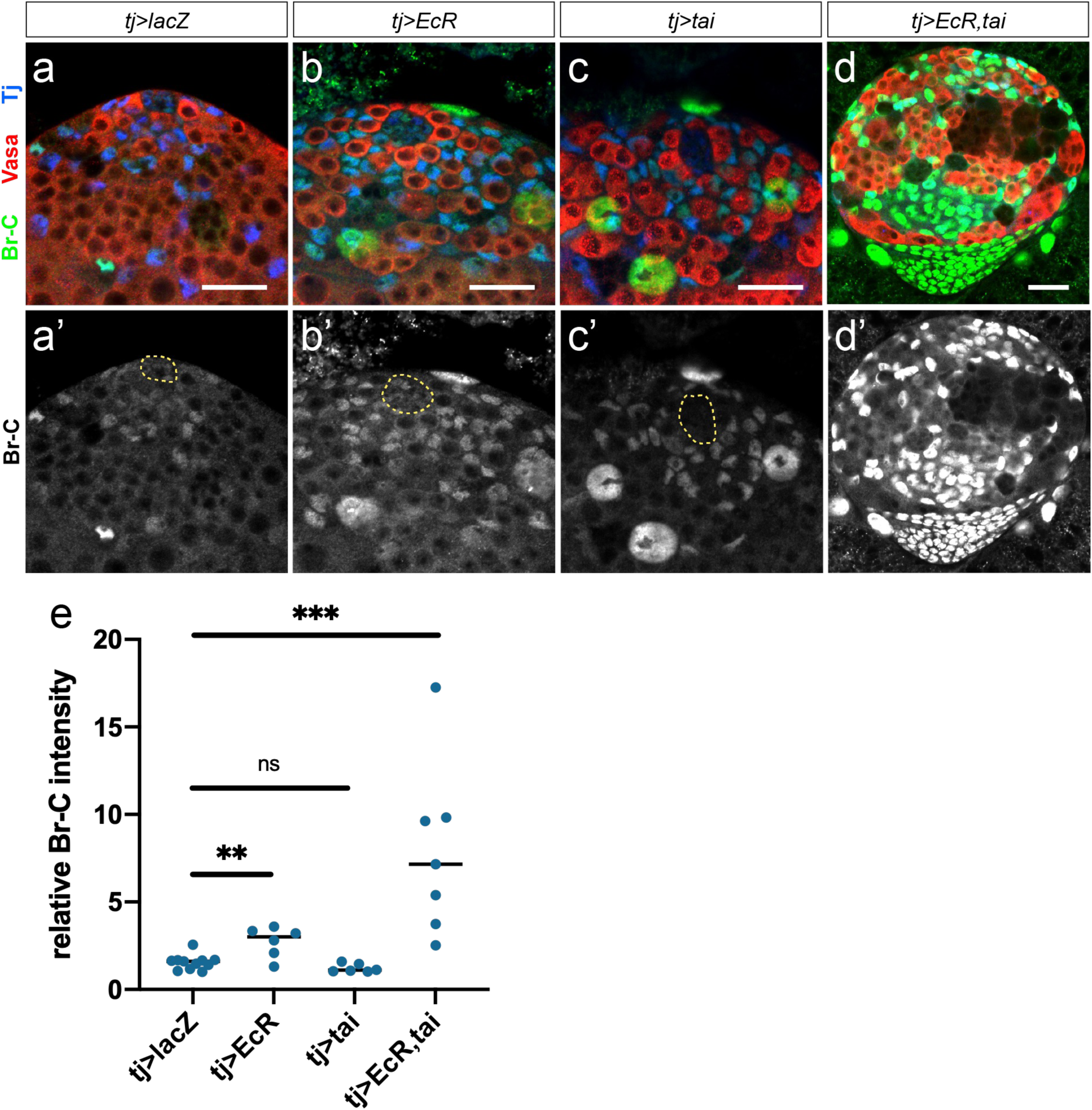
(a-d) Br-C expression in LL3 control testes upon somatic over-expression of EcR (b), Tai (c), or EcR and Tai simultaneously (d). (e) Quantification of relative Br-C intensity from a-d. Scale bars = 25 μm.

**Extended Data Figure 6.**
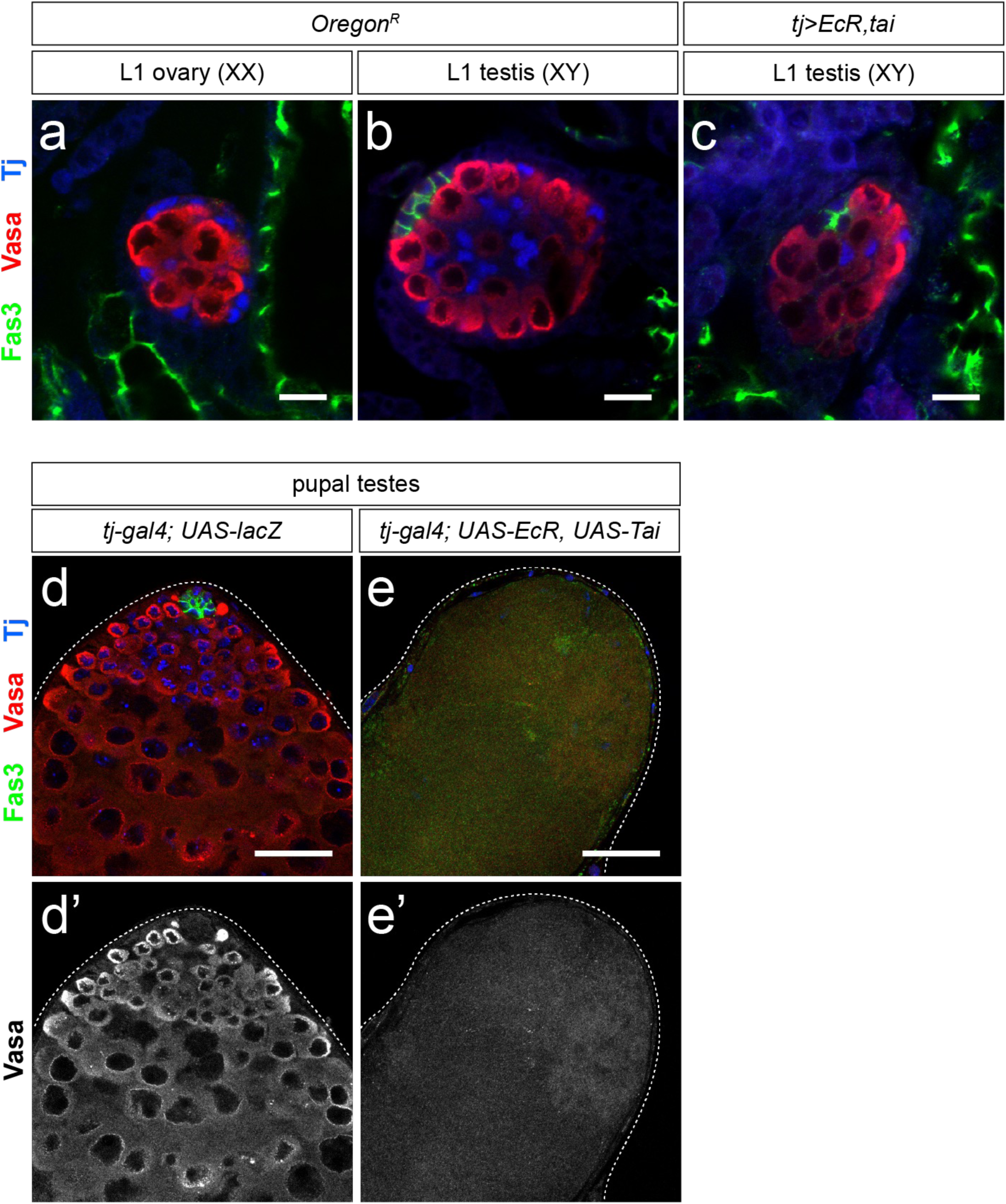
(a-c) Representative images of early L1 (24-28h after egg lay, AEL) gonads, identified by the presence of a Fas3-positive hub (green). Ovaries lack a hub (a) while testes contain a Fas3-positive hub (green) by this stage (b). III staining. Co-expression of *EcR* and *tai* in the early somatic gonad (c) leads to a visibly smaller hub that contains significantly fewer hub cells than control L1 testes (see Fig. 3e for quantification). Vasa (red) labels the germline, Tj (blue) labels somatic gonadal precursor cells, which give rise to the somatic gonad in both sexes. (d-e) Pupal testes upon EcR/Tai co-expression. In a control pupal testis (d), the germline and somatic stem cells are maintained by the presence of a hub (Fas3, green) and thus differentiating germ cells (Vasa, red) are observed in the testis. In some pupal testes, following EcR/Tai co-expression in the somatic gonad (e), no Fas3-positive hub cells are present and the absence of Vasa and DAPI staining indicates that both germline and somatic cell lineages have been lost. DAPI-positive muscle sheath cells can be found at the periphery of the testis. Scale bars = 10 μm.

**Extended Data Figure 7.**
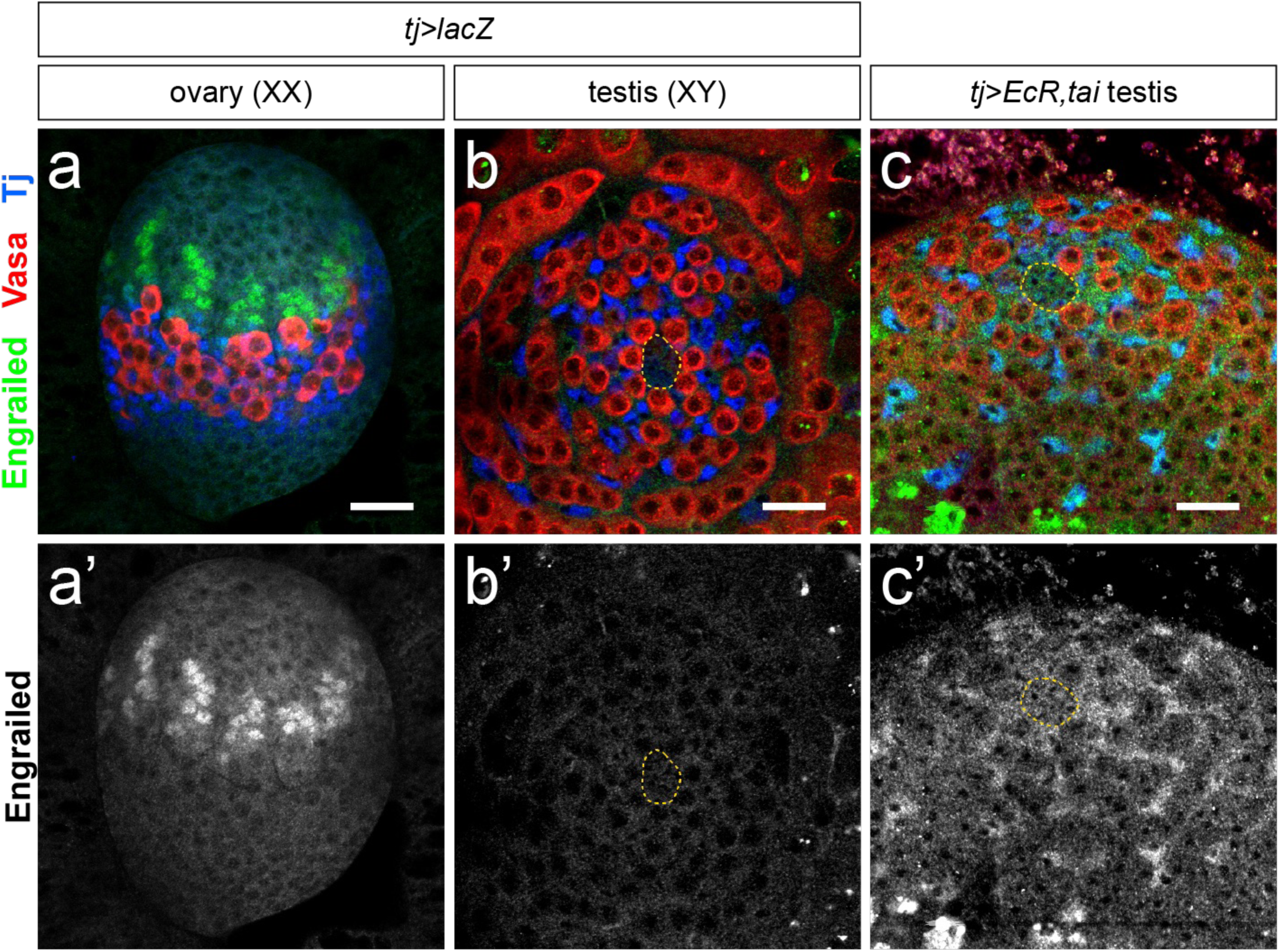
(a-c) Expression of the TF marker Engrailed (En) in LL3 gonads. En (green) labels TF cells in a control ovary (a) but is not detected in the niche of a control testis (b). En is not detected in hub cell nuclei upon EcR/Tai co-expression in the somatic gonad (c). En induction is observed in the cyst cell lineage upon EcR activation (c). In b and c, the hub is outlined with a yellow dotted line. Scale bars = 25 μm.

**Extended Data Figure 8.**
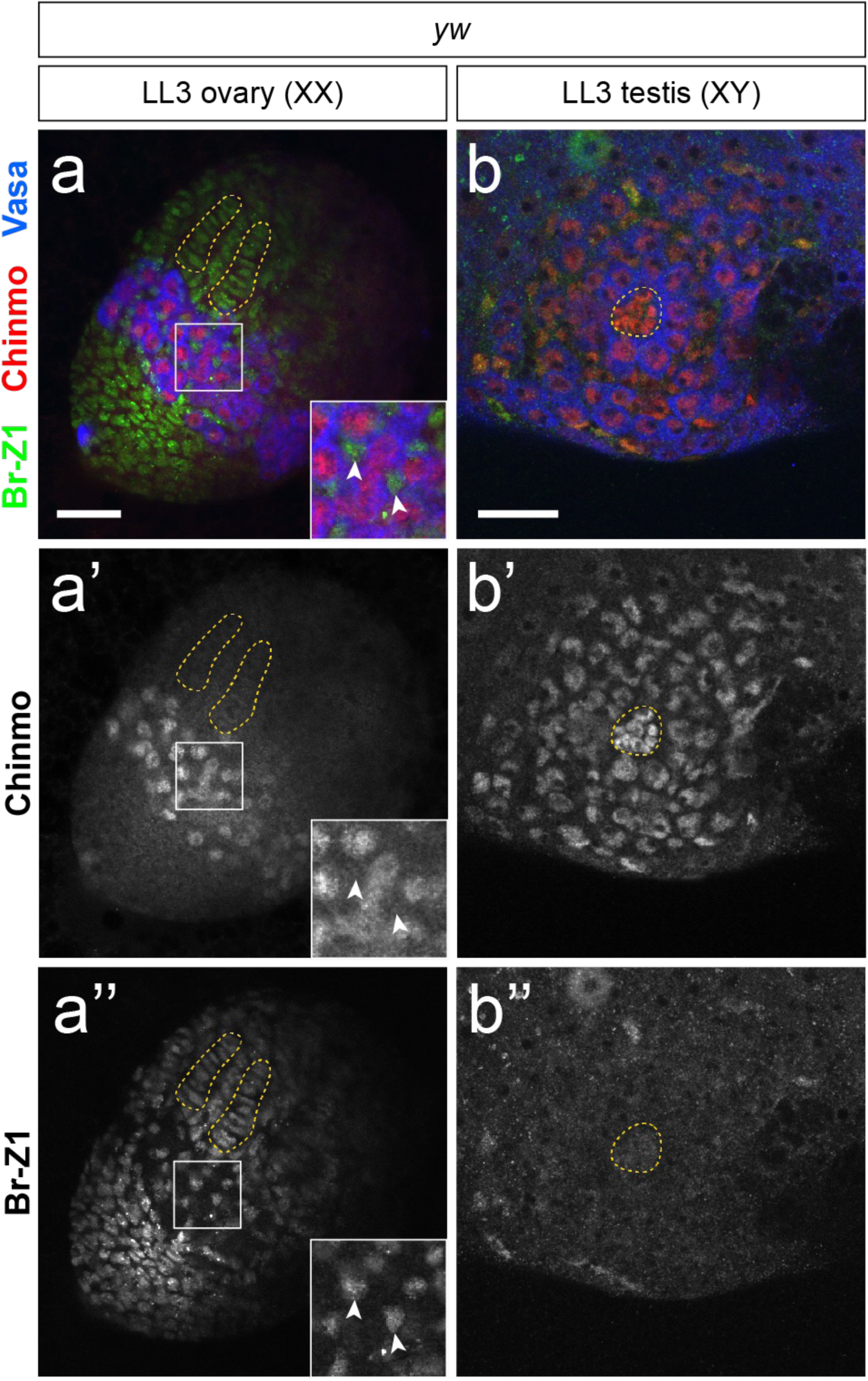
Mutually exclusive expression of Chinmo (red) and Br-Z1 (green) in somatic cells of LL3 gonads. In an LL3 ovary (a), Chinmo is absent from somatic cells (a’, inset arrowheads) while Br-Z1 is expressed broadly in the female somatic gonad. We note that Chinmo is expressed in female larval progenitor germ cells (PGCs) (a’). In an LL3 testis (b), Chinmo is expressed in the hub, CySCs/cyst cells, and GSCs/early germ cells. Br-Z1 is not detectable in the hub or early cyst cells. Vasa (blue) labels the germline. Scale bars = 25 μm.

**Extended Data Figure 9.**
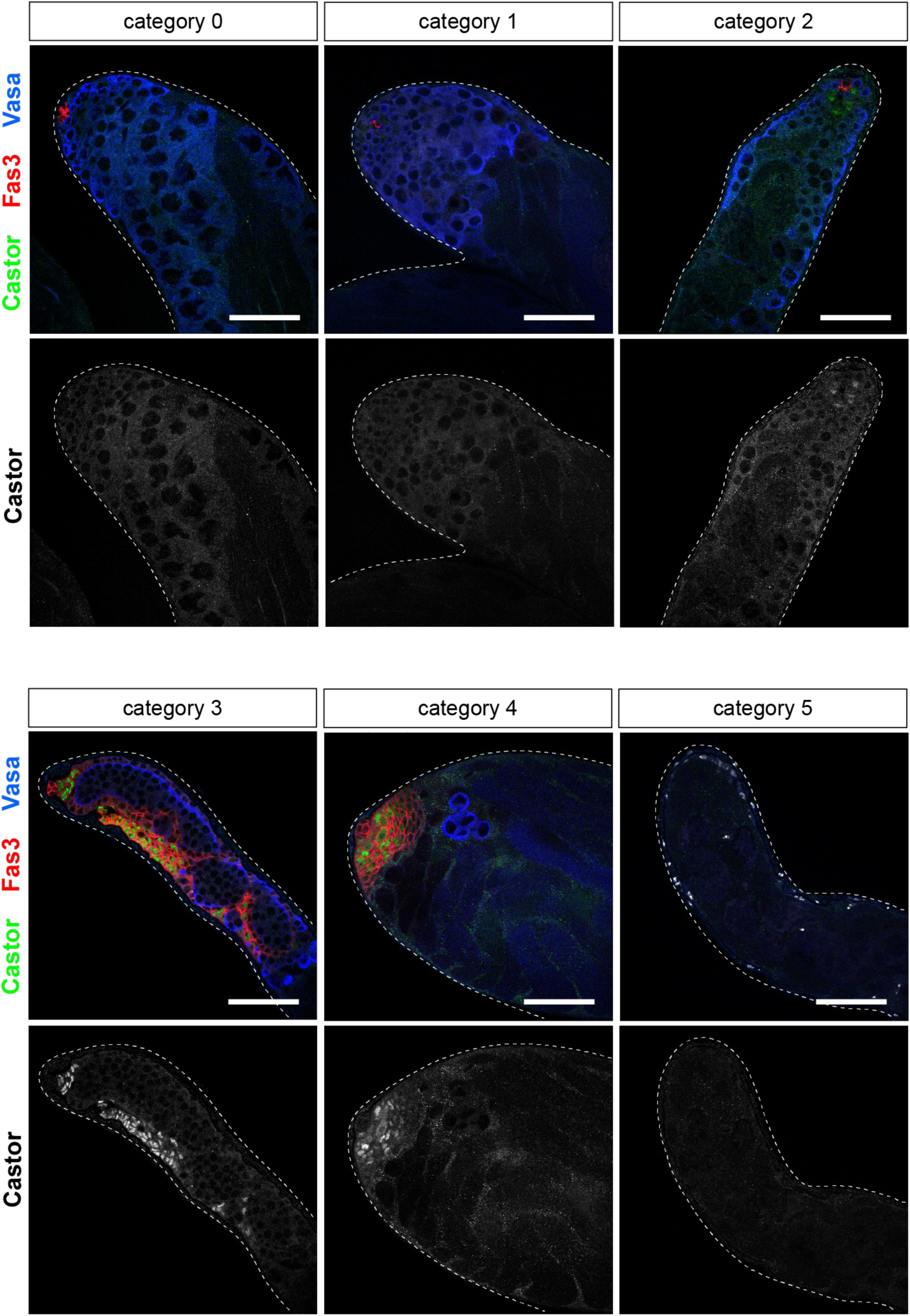
Phenotypic category descriptions and representative images for rescue quantification in Figure 4i. (a) Category 0: no evidence of feminization (accumulation of Fas3/Castor-expressing somatic aggregates), early/late spermatogonia and large-nuclei spermatocytes present. (b) Category 1: No evidence of feminization, but germline defects seen (shortening of mitotic region and/or absence of large-nuclei spermatocytes). (c) Category 2: mild feminization (a few cells express Fas3 and/or Castor, but aggregates have not yet formed). (d) Category 3: moderate feminization (follicle-like aggregates present clearly expressing Fas3 and Castor, few germ cells are present and mostly seem arrested in spermatogonial stage). (e) Category 4: severe feminization (large follicle-like aggregates present, few/no germ cells remain). (f) Category 5: acellular. Vasa (blue) labels the germline. Scale bars = 50 μm.

